# Oxytocin neurons in the paraventricular and supraoptic hypothalamic nuclei bidirectionally modulate food intake

**DOI:** 10.1101/2024.11.20.624599

**Authors:** Jessica J. Rea, Clarissa M. Liu, Anna M.R. Hayes, Rita Ohan, Grace M. Schwartz, Alexander G. Bashaw, Molly E. Klug, Lea Decarie-Spain, Yedam Park, Alicia E. Kao, Valery Grinevich, Scott E. Kanoski

## Abstract

Oxytocin (OT) is a neuropeptide produced in the paraventricular (PVH) and supraoptic (SON) nuclei of the hypothalamus. Either peripheral or central administration of OT suppresses food intake through reductions in meal size. However, pharmacological approaches do not differentiate whether observed effects are mediated by OT neurons located in the PVH or in the SON. To address this, we targeted OT neuron-specific designer receptors exclusively activated by designer drugs (DREADDs) in either the PVH or SON in rats, thus allowing for evaluation of food intake following selective activation of OT neurons separately in each nucleus. Results revealed that DREADDs-mediated excitation of PVH OT neurons reduced consumption of both standard chow and a high fat high sugar diet (HFHS) via reductions in meal size. On the contrary, SON OT neuron activation had the opposite effect by increasing both standard chow and liquid sucrose consumption, with the former effect mediated by an increase in meal size. To further examine the physiological role of OT neurons in eating behavior, a viral-mediated approach was used to silence synaptic transmission of OT neurons separately in either the PVH or SON. Results from these studies revealed that PVH OT neuron silencing significantly increased consumption of HFHS by increasing meal size whereas SON OT neuron silencing reduced chow consumption by decreasing meal size. Collectively these data reveal that PVH and SON OT neurons differentially modulate food intake by either increasing or decreasing satiation signaling, respectively.

## INTRODUCTION

Oxytocin (OT) is a nine amino-acid neuropeptide produced predominantly within the paraventricular hypothalamic nucleus (PVH) and supraoptic nucleus (SON) of the hypothalamus [1–3]. OT mediates a diverse set of functions both through its actions following release from the posterior pituitary into the periphery as a hormone, as well as through neuropeptide OT signaling within the central nervous system where oxytocin receptors (OTRs) are expressed widely throughout the brain [4]. OT acts on a G-protein coupled OTR to impact a wide range of behaviors, including social behavior, reproduction, stress, and food intake control [5–7]. With regards to its function in regulating appetite and food consumption, OT has been shown to be a potent anorexigenic signal that reduces food intake across species when administered both centrally and in the periphery [8–11]. Pharmacological studies have identified OT’s reduction of food intake to be driven by its role in boosting satiation signals coming into the hindbrain from the gut, leading to earlier meal termination and reduced meal size [12; 13]. Studies in both rodents and humans have demonstrated that these effects are even stronger in overweight and obese individuals [14–16]. For these reasons OT has been identified as an exciting potential therapy for the treatment of obesity [17–19].

Our knowledge of the role of OT in appetite and food intake control comes predominantly from pharmacological studies that utilize the administration of exogenous OT centrally or into the periphery [8; 11; 12; 15; 16]. While these approaches are clinically relevant and are informative for understanding the role of specific populations of OTRs in eating behavior, pharmacological studies do not offer insights into the function of hypothalamic OT neurons that release OT along with other neurotransmitters. This is a critical gap as evidence has shown that PVH and SON OT neuron subpopulations have different morphological compositions, different OT and neurotransmitter release dynamics, and these subpopulations have both distinct and overlapping monosynaptic inputs and projections [20; 21]. Given that the brainstem is a potent site of action for OT pharmacology-mediated meal size reduction [12; 13; 22], and that PVH but not SON OT neurons project directly to the brainstem [20], we hypothesized that PVH and SON neurons may play distinct roles in modulating feeding behavior, with the PVH subpopulation yielding more potent anorexigenic action than the SON subpopulation. Here we utilized both gain-of-function (viral-mediated chemogenetic activation) and loss-of-function (synaptic transmission silencing) approaches to investigate the role of the PVH and SON OT neuron subpopulations, separately and together, in the control of feeding behavior. Given that OTR pharmacological studies identify macronutrient-dependent responses [23–25], we examined effects on consumption of standard chow, a high-fat and high-sugar diet, and liquid sucrose separately.

## MATERIALS & METHODS

### Animals

Adult male Sprague-Dawley rats (Envigo, Indianapolis, IN; PND >60; 320-450 g at the start of the experiment) were individually housed in a temperature-controlled vivarium with *ad libitum* access (except where noted) to water and food (LabDiet 5001, LabDiet, St. Louis, MO) on a 12h:12h reverse light/dark cycle. All procedures were approved by the Institute of Animal Care and Use Committee at the University of Southern California.

### Surgery

For all surgical procedures, rats were anesthetized and sedated via intramuscular injections of ketamine (90 mg/kg), xylazine (2.8 mg/kg), and acepromazine (0.72 mg/kg). Rats were also given analgesic (subcutaneous injection of ketoprofen [5 mg/kg]) after surgery and once daily for 3 subsequent days thereafter. All rats recovered for at least one-week post-surgery prior to experimental procedures.

### Intracranial cannula implantation for drug delivery

For pharmacological delivery into the lateral ventricle (LV) for intracerebroventricular (ICV) drug (clozapine N-oxide [CNO]) administration, rats were surgically implanted with a unilateral indwelling guide cannula (26-gauge, Plastics One, Roanoke, VA) using the stereotaxic coordinates, relative to the location of bregma: -0.90 mm anterior/posterior (AP), +1.80 mm medial/lateral (ML), and -2.60 mm dorsal/ventral (DV) with the DV coordinate zeroed at the surface of the skull before being lowered into the brain. Cannula were affixed to the skull as previously described using jeweler’s screws and dental cement [26]. Following a week of recovery, the optimal injector tip length for infusion into the LV was determined by injecting 5-Thio-D-Glucose (5-TG) (Sigma-Aldrich, St. Louis MO) into the LV and measuring change in blood glucose. Rats were mildly food restricted before 5-TG testing [27]. An initial blood glucose reading was taken an hour after the dark cycle onset using a OneTouch monitor with blood taken from the tip of the tail from a small cut by a sterile razor blade. Starting with an injector that extends 2.0 mm beyond the end of the guide cannula, 2 uL of a 105 mg/mL 5-TG solution was then infused into the LV using a Hamilton microinjector.

Blood glucose readings were taken 30 min and 1 hr following infusion of 5-TG. An animal was considered to pass with a given tip length targeting the ventricle if their blood glucose approximately doubled in that time. The procedure was repeated on subsequent days as needed for an animal with injector tip lengths of 2.50 mm and 3.00 mm until the animal passed. This passing tip length was then used for all subsequent ICV drug deliveries.

### Intracranial virus injection

Stereotaxic injections of viruses were delivered using a micro-infusion pump (Harvard Apparatus, Cambridge, MA, USA) connected to a 26-gauge microsyringe injector attached to a PE20 catheter and 10 µL Hamilton syringe. The flow rate was calibrated and set to 5 µL/min. The volume of each viral infusion was 300 nL and injectors were left in place for 2 min post-infusion. Following viral infusions, animals were either implanted with indwelling LV ICV cannula as described above, or surgically closed with sutures. Experiments occurred starting 3 weeks after viral injection to allow for virus transduction and expression. Successful expression was confirmed postmortem in animals via immunohistochemistry (IHC) staining as described below. All OTp-driven viruses were produced and validated by Dr. Valery Grinevich’s laboratory at the Central Institute of Mental Health, University of Heidelberg, Germany.

For chemogenetic activation of OT neurons via DREADDs (Figure 1A) an AAV1/2_OTp_hM3D(Gq)-mcherry (OTp DREADDs; [28]) was bilaterally injected to target regions at the following coordinates: paraventricular hypothalamus (PVH) -1.80 mm AP, +/-0.35 mm ML, -8.0 mm DV; and supraoptic nuclei (SON) -1.10 mm AP, +/- 1.80 mm ML, -9.20 mm DV (0 reference point for AP, ML and DV at bregma). Representative photomicrographs of OTp DREADDs viral expression in each region are shown in Figure 1B.

**Figure 1.**
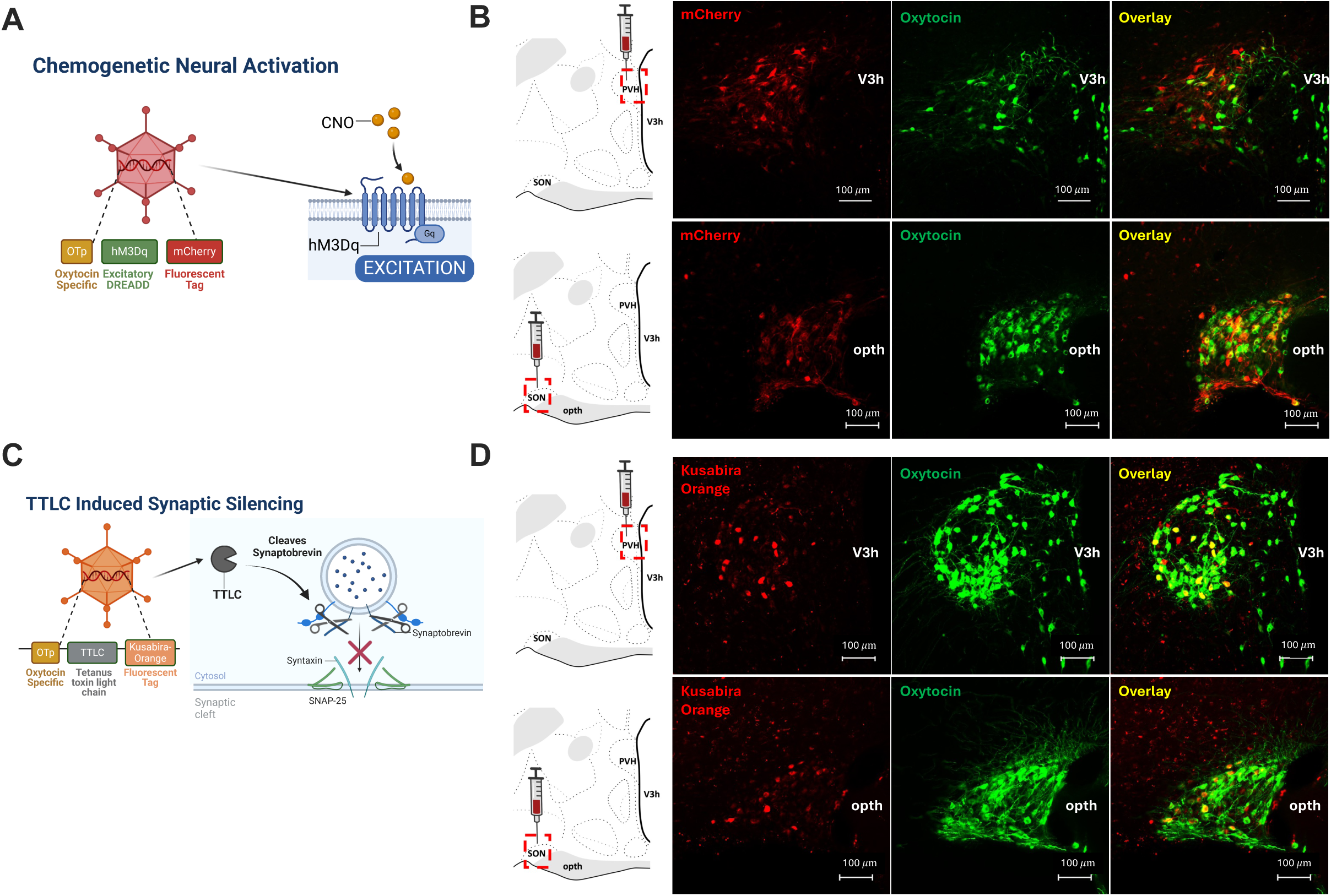
Viral mediated approaches to investigate effects of OT neuron activation and silencing. **(A)** Diagram depicting viral vector composition for the expression of excitatory hM3Dq DREADDs in OT neurons (OTp DREADDs). **(B)** Representative photomicrographs showing viral-induced expression of mcherry in OT neurons confirmed with immunohistochemistry following injection of OTp DREADDs into the PVH and SON, respectively. **(C)** Diagram depicting viral approach to silence synaptic transmission through the expression of tetanus toxin light chain (TTLC) to cleave synaptobrevin and prevent vesicular fusion in OT neurons (OTp TTLC). **(D)** Representative photomicrographs showing viral-induced expression of koomassi orange (ko) in OT neurons confirmed with immunohistochemistry following injection of OTp TTLC into the PVH and SON respectively. (Abbreviations: CNO = clozapine-N-oxide V3h = 3^rd^ ventricle, opth = optic chiasm).

Synaptic silencing of vesicular release of OT was accomplished using AAV1/2_OTp_TTLC-ko_WPRE (OTp TTLC; [29]) virus to express the tetanus-toxin light chain (TTLC) protein which cleaves vesicle-associated membrane protein synaptobrevin to prevent vesicle fusion and release (Figure 1C). Additionally, this virus expresses a koomassi orange (ko) fluorescent reporter protein that can be visualized under a fluorescent microscope. This virus was targeted to PVH and SON regions using the same coordinates listed above. Representative photomicrographs of OTp TTLC viral expression in each region is shown in Figure 1D.

To control for effects of viral infusions, we used an OT promoter virus absent DREADDs or TTLC constructs (AAV1/2_OTp_Venus; [30]) as a control virus which drives the expression of the yellow fluorescent protein Venus selectively in OT neurons [31].

### Immunohistochemistry (IHC) and microscopy

Rats were anesthetized and sedated via intramuscular injections of ketamine (90.1 mg/kg)/xylazine (2.8 mg/kg)/acepromazine (0.72 mg/kg) cocktail, then transcardially perfused with 0.9% sterile saline (pH 7.4) followed by 4% paraformaldehyde (PFA) in 0.1M borate buffer (pH 9.5). Brains were dissected out and post-fixed in PFA with 12% sucrose for 24 h, then flash-frozen in isopentane cooled in dry ice. Brains were sectioned to 30-μm thickness on a freezing microtome and sections were collected and stored in antifreeze solution at −20°C until further processing.

The following IHC fluorescence labeling procedures were adapted from previous work [32]. Guinea pig anti-Oxytocin-neurophysin 1 (1:10,000 dilution; Abcam, Cambridge, United Kingdom; Catalog #:ab228508, Lot #:GR3357270-1; Clonality: Polyclonal) and rabbit anti-RFP (1:2000 dilution, Rockland Inc., Limerick, PA, USA; Catalog #:600-401-379; Clonality: Polyclonal) were the two antibodies used. Antibodies were prepared in 0.02M potassium phosphate-buffered saline (KPBS) solution containing 0.2% sodium azide and 2.0% normal donkey serum and stored at 4°C overnight. After a series of six washes with 0.02M KPBS, brain sections were incubated in a secondary antibody solution. The two secondary antibodies used, donkey anti-rabbit Cy^TM^3 (Catalog #:711-165-152) and donkey anti-guinea pig Alexa Fluor (AF) 488 (Catalog #:706-545-148) had a 1:500 dilution and were stored overnight at 4 °C (Jackson ImmunoResearch; West Grove, PA, USA). Sections were then mounted and cover-slipped with using 50% glycerol in 0.02M KPBS and clear nail polish was used to seal the coverslip onto the slide.

Antibody tagging of OT first involved washing the brain sections on a motorized rotating platform in the following order (overnight incubations on a motorized rotating platform at 4°C): (1) 0.02M KPBS (changing KPBS every 5 min for 30 min), (2) 0.3% Triton X-100 in KPBS (30 min), (3) KPBS (changing KPBS every 5 min for 15 min), (4) 2% donkey serum in KPBS (10 min), (5) 2% normal donkey serum, 0.2% sodium azide, and guinea pig anti-OT [1:10,000; guinea pig anti-OT] and rabbit anti-RFP [1:2000; rabbit anti-RFP] in KPBS (24 h), (6) KPBS (changing KPBS every 10 min for 1 h), (7) 2% normal donkey serum, 0.2% sodium azide and secondary antibodies (1:500; donkey anti-guinea pig AF488 and donkey anti-rabbit Cy3; overnight) in KPBS (30 h), (8) KPBS (changing KPBS every 2 min for 4 min). Photomicrographs were acquired using a Nikon 80i (Nikon DS-QI1,1280 x 1024 resolution, 1.45 megapixel) microscope under epifluorescence.

### Characterization of OT DREADDs and TTLC expression

For OT DREADDs experiments, immunostaining for RFP to amplify the mCherry signal and label OT neurons was conducted as described above. Counts were performed in sections from the Swanson Brain Atlas levels 22-27, which encompasses OT-containing neurons in the PVH and SON, for colocalization of OT and fluorescence reporter for OT DREADDs. For OT DREADDs experiments, animals were excluded from analysis if fewer than 40% of the total number of OT neurons were transduced with RFP (based on IHC staining for OT). All animals that met this criterion were included for experimental analysis. The percentage of OT that express the viral DREADDs marker (# expressing RFP/# total immunoreactive OT neurons) for the PVH was: Mean = 0.7603, SEM = 0.0331, SD = 0.119. For the SON, the percentage was: Mean = 0.7262, SEM = 0.0143, SD = 0.0757.

For OT neuron synaptic silencing (TTLC) experiments IHC was performed as described above to label OT neurons using a green fluorescent protein. Counts were performed in sections from the Swanson Brain Atlas levels 22-27, which encompasses OT-containing neurons in the PVH and SON, for colocalization of OT neurons and fluorescence reporter for OT TTLC virus which consisted of a kusabira orange (KO) protein that could be visualized using the TRITC filter on the microscope. For OT neuron silencing experiments, animals were excluded from analysis if fewer than 40% of the total number of OT neurons were transduced with kusabira orange. All animals that met this criterion were included for experimental analysis. Experimenters performing the counting were blind to experimental assignments. The percentage of OT that express the viral TTLC marker (# expressing KO/# total immunoreactive OT neurons) for the PVH was: Mean = 0.598, SEM = 0.081, SD = 0.244. For the SON, the percentage was Mean = 0.7644, SEM = 0.044, SD = 0.1538.

### Drug preparation

For chemogenetic activation of OT neurons, 2 µL (18 mM/2 µL) CNO or vehicle daCSF (33% DMSO in 66% aCSF) is administered ICV through a microsyringe connected to an injector tip. Injector tip was left in place for 30 seconds before removal and any instances of backflow from the cannula upon injector removal were noted. Doses of CNO and vehicle were based on previous work [32; 33]. Prior to drug injections animals were handled and prepared for injections. Injections of CNO or vehicle occurred 45-60 min prior to the start of measuring feeding behavior.

### Caloric intake studies

Animals were housed in a reverse light/dark cycle (lights off at 11:00 a.m.). Home cages were integrated in a BioDAQ food monitoring intake system to which they were habituated to for at least 5 days prior to any treatments. For pharmacological experiments on test days, home cage food access was removed 1 h prior to dark onset. For chemogenetic activation of OT neurons, drug treatments were counterbalanced across animals using a within-subject design. Infusions of CNO or daCSF occurred 45-60 min prior to the start of the dark cycle, at which point access to food was restored and food intake parameters were recorded for the subsequent 6 h. Drug treatments were separated by 2 days.

Chow experiments were conducted with animals maintained on ad libitum chow (LabDiet 5001, LabDiet, St. Louis, MO) housed within the BioDAQ system. Chemogenetic activation experiments consisted of two treatment days separated by two intervening washout days where chow intake was recorded from the onset of the dark cycle. For chronic silencing experiments animals were maintained on chow for a period of habituation as well as for recording 5 consecutive days of meal patterns starting at the onset of the dark cycle that were averaged together. High fat high sugar diet (HFHS) experiments were conducted using a 45 kcal% fat diet (Research Diets Inc. New Brunswick, NJ; Catalog #: D12451) which the animals were habituated to and maintained on for at least 5 days prior to test days or recording. For sucrose experiments animals received two habituation days to a 11% weight by volume sucrose solution (colored with blue food dye) with 2 h access to sucrose solution in their home cage using a liquid BioDAQ monitor. Habituation was followed by two test days that recorded sucrose intake for 2 h starting at the onset of the dark cycle, during which time regular food was not available.

For analysis of BioDAQ data for chow and HFHS conditions an inter-meal interval (IMI) of 900 s was used to separate individual eating bouts into separate meals. Data was binned into 12 bins of 30 min each which were used to calculate values for each of the time points depicted for DREADDs experiments (0.5 h, 1 h, 2 h, 4 h, 6 h). Intake parameters were only evaluated for up to 6 h for DREADDs experiments based on the half-life of CNO, as well as previous results from our lab in which CNO-mediated effects on food intake did not extend beyond 6 h using a similar food intake experimental design [32–34]. For chronic silencing experiments data was extracted for each day of a 5-day period and then averaged together to get final values. Sucrose tests were analyzed with a 300 sec (5 min) IMI to separate liquid sucrose meals [35], and the test only occurred for 2 h total length with data binned into 30 min intervals.

### Statistical analysis

Statistical analyses were performed using GraphPad Prism 10.3.1 software (GraphPad Software Inc., San Diego, CA, USA). Data are expressed as mean +/- SEM. Statistical test details can be found in the figure legends and “*n*’s” refers to the number of animals for each condition. Significance was considered at p<0.05.

## RESULTS

### Oxytocin neurons in the PVH reduce food intake via meal size reduction

To activate OT neurons, an excitatory DREADDs virus was infused into the PVH region (Figure 1A). Viral expression was confirmed through IHC (as described above) and visualized under fluorescent microscopy with a representative photomicrograph shown in Figure 1B. CNO-mediated activation of PVH OT neurons significantly reduced cumulative intake of standard chow at the 0.5-h, 1-h, and 6-h time points (Figure 2A), an effect driven by a decrease in average meal size (Figure 2C) as there was no effect on meal frequency (Figure 2B). In animals maintained on 45% kcal from fat HFHS, activation of PVH OT neurons with CNO on test days resulted in a trend towards a decrease in cumulative HFHS intake at 4 h (p<0.07; Figure 2D). Infusion of CNO had no effect on HFHS meal frequency (Figure 2E), but did significantly reduce average HFHS meal size at 4 h (Figure 2F). In a 2-h consumption test with access to an 11% sucrose solution without chow available, activation of PVH OT neurons had no effect on cumulative sucrose intake (Figure 2G) or average liquid sucrose meal size (Figure 2I). There was a reduction in liquid sucrose meal frequency at the 2-h time point with the infusion of CNO vs. vehicle (Figure 2H). However, this effect was not sufficient to reduce overall calories of sucrose consumed.

**Figure 2.**
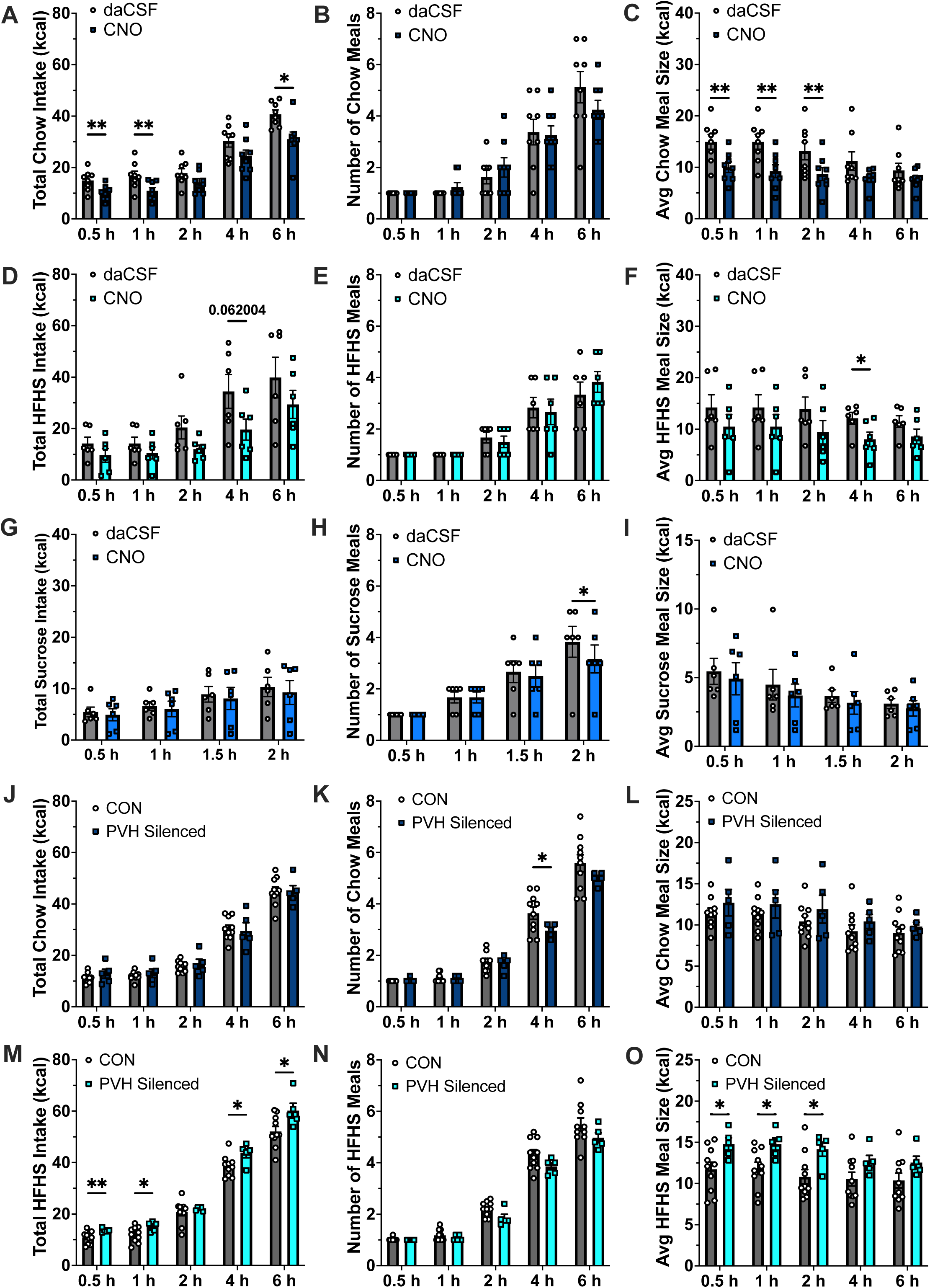
Chemogenetic activation of PVH OT neurons reduces chow intake, while silencing of PVH OT neurons increases HFHS intake. **(A-C)** Activation of PVH OT neurons via ICV CNO administration reduced cumulative chow intake at 0.5 h (t(7)=3.691, p=0.008), 1 h (t(7)=3.940, p=0.006), and 6 h (t(7)=2.942, p=0.021), an effect that was driven by a decrease in average chow meal size (0.5 h (t(7)=3.669, p=0.008), 1 h (t(7)=4.310, p=0.004), and 2 h (t(7)=3.667, p=0.008)) without affecting meal frequency for chow intake. **(D)** Activation of PVH OT neurons resulted in a trend towards decreased cumulative HFHS intake at 4 h (t(5)=2.385, p=0.062). **(E)** There was no effect of PVH OT neuron activation on HFHS meal frequency, however **(F)** there was a significant decrease in average HFHS meal size at 4 h (t(5)=3.529, p=o.017). **(G-I)** Activation of PVH OT neurons had no effect on cumulative sucrose intake or average liquid sucrose meal size, though it did significantly decrease 2 h liquid sucrose meal frequency (t(5)=3.162, p=0.025). **(J-L)** Silencing synaptic release of PVH OT neurons had no effect on average daily intake of chow or average chow meal size, though the silencing did cause a significant reduction in meal frequency 4 h into the dark cycle (Fisher’s LSD planned comparisons 4 hr: t(12.22)=2.548, p=0.025 CON vs PVH Silenced). **(M)** Silencing of PVH OT neurons significantly increased average cumulative HFHS intake at all time points but 2 h (Fisher’s LSD planned comparisons 0.5 h: t(12.02)=3.249, p=0.007; 1 h: t(12.19)=2.437, p=0.031; 4 h: t(8.369)=2.311, p=0.0482; 6 h: t(7.785)=2.320, p=0.0498; 12 h: t(12.86)=2.232, p=0.044 and 24 h: t(12.17)=2.192, p=0.0485). **(N)** There was no difference between CON and PVH Silenced groups in HFHS meal frequency. **(O)** PVH OT neuron silencing increased average HFHS meal size at 0.5 h, 1 h, 2 h, and 24 h from dark onset (Fisher’s LSD planned comparisons 0.5 h: t(12.25)=2.668, p=0.02; 1 h: t(11.24)=2.781, p=0.0175; 2 h: t(11.27)=2.723, p=0.0194; and 24 h: t(9.530)=2.395, p=0.0388). (DREADDs activation: chow n=8; sucrose n=6; HFHS n=6; within-subjects design for drug treatments; TTLC Silencing: CON n=10; PVH Silenced n=5, between-subjects for group; Data are means ± SEM; *p<0.05, **p<0.01).

To chronically silence PVH OT neuron synaptic transmission for loss-of-function analyses, a TTLC expressing virus was infused into the PVH region (Figure 1C); representative photomicrographs of viral expression are visualized in Figure 1D. Chronic silencing of synaptic release from PVH OT neurons had no effect on average daily cumulative chow intake (Figure 2J) or average chow meal size (Figure 2L), however it did reduce average chow meal frequency 4 h into the nocturnal/dark cycle (Figure 2K). When placed on a HFHS diet, chronic silencing of PVH OT neurons increased average daily cumulative HFHS intake at 0.5 h, 1 h, 4 h and 6 h into the nocturnal cycle (Figure 2M), an effect driven by an increase in average HFHS meal size at the 0.5 h, 1 h, and 2 h time points (Figure 2O), with no change in HFHS meal frequencies (Figure 2N). Despite these effects on early nocturnal phase food intake, there were no significant differences in PVH OT neuron silenced animals and controls for bodyweight (Supplemental Figure 1A) or change in body weight since surgery (Supplemental Figure 1B). Taken together with results from excitatory DREADDs experiments above, these findings suggests that the endogenous role of PVH OT neurons involves decreasing food intake by reducing meal size, particularly during periods of active nocturnal feeding.

### Oxytocin neurons in the SON increase food intake through elevated meal size

The same viral mediated approach to express excitatory DREADDs was used to target OT neurons in the SON (Figure 1A), with representative photomicrographs of viral expression shown in Figure 1B. Unexpectedly, LV infusion of CNO to activate SON OT neurons increased cumulative chow intake at the 0.5-h and 1-h time points (Figure 3A). This hyperphagic effect was primarily driven by a significant increase in average chow meal size (significant at the 0.5-, 1-, and 4-h time points; Figure 3C) as there was no effect on meal frequency (Figure 3B). In animals maintained on the HFHS, activation of SON OT neurons had no effect on cumulative HFHS intake (Figure 3D) or average HFHS meal size (Figure 3F) but did result in a significant decrease in HFHS meal frequency at 6 h (Figure 3E). In a 2-h consumption test with access to an 11% sucrose solution without chow available, activation of SON OT neurons significantly increased cumulative sucrose intake at the 0.5-h, 1-h, and 1.5-h time points (Figure 3G). There was no significant difference in average liquid sucrose meal size (Figure 3I), but there was a significant increase in liquid sucrose meal frequency at 1.5 h and at 2 h (Figure 3H).

**Figure 3.**
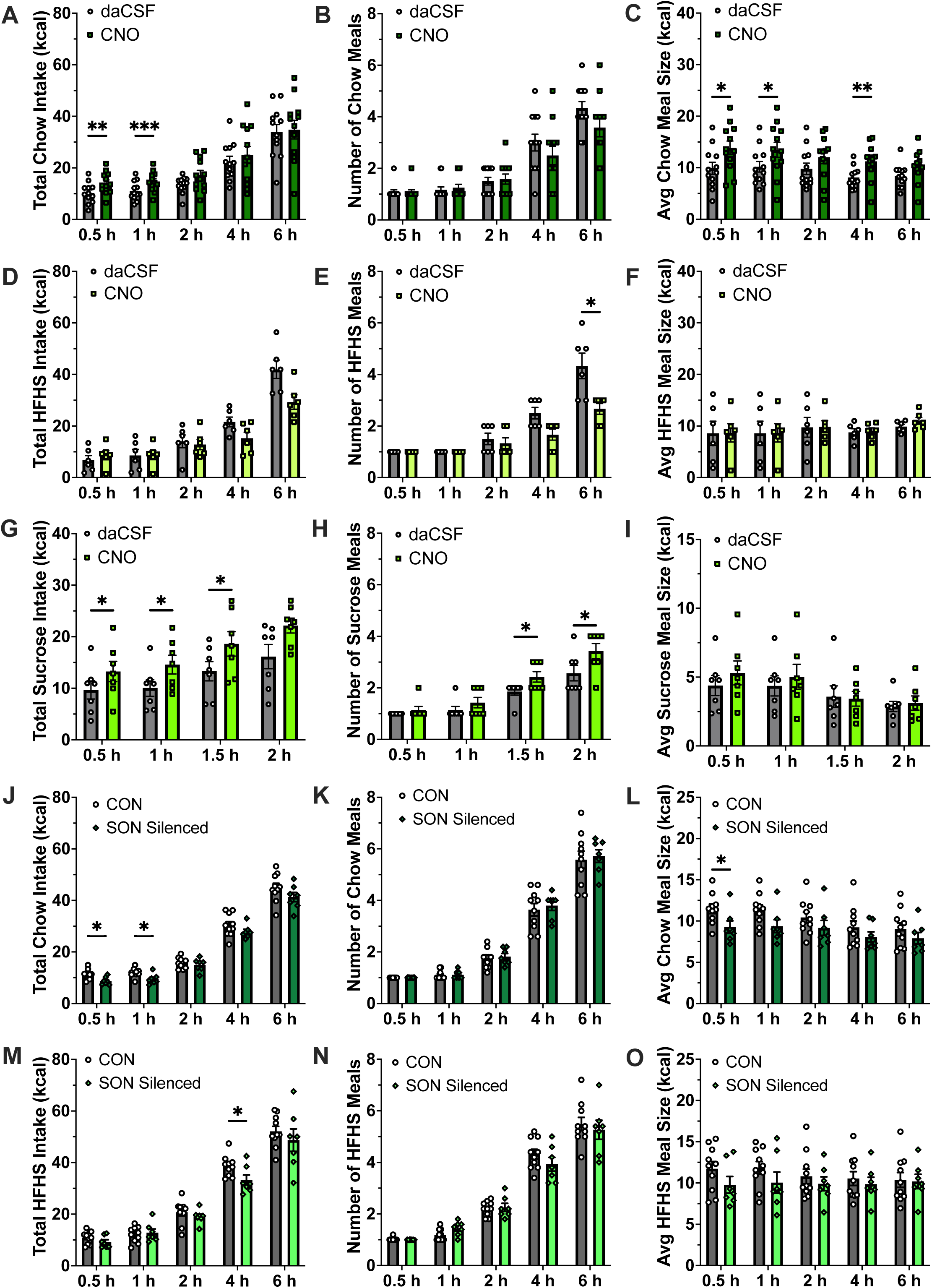
Chemogenetic activation of SON OT neurons increases chow and sucrose intake while silencing decreases chow intake. **(A-C)** Activation of SON OT neurons via ICV CNO administration increased cumulative chow intake at 0.5 h (t(11)=3.678, p=0.004), and 1 h (t(11)=4.475, p=0.0009), an effect that was driven by an increase in average chow meal size (0.5 h (t(11)=3.001, p=0.012), 1 h (t(11)=2.509, p=0.029), and 4 h (t(11)=3.569, p=0.0044)) without affecting meal frequency for chow intake. **(D-F)** There was no effect of SON activation on cumulative HFHS intake or average HFHS meal size, though there was a significant decrease in HFHS meal frequency at 6 h (t(5)=2.712, p=0.042). **(G)** Activation of SON OT neurons significantly increased cumulative sucrose intake at 0.5 h (t(6)=2.868, p=0.029), 1 h (t(6)=3.102, p=0.021), and 1.5 h (t(6)=2.449, p=0.0498). **(H)** Activation of SON OT neurons resulted in an increase in liquid sucrose meal frequency at 1.5 h (t(6)=2.828, p=0.03), and 2 h (t(6)=2.452, p=0.045), while **(I)** there was no effect on average liquid sucrose meal size. **(J-L)** Silencing synaptic release from SON OT neurons significantly decreased average cumulative chow intake at 0.5 h and 1 h from dark onset (Fisher’s LSD planned comparisons 0.5 h: t(14.68)=2.944, p=0.01; 1 h: t(11.61)=2.748, p=0.018), an effect driven by a decrease in 0.5 hr chow meal size (Fisher’s LSD planned comparisons 0.5 h: t(12.14)=2.327, p=0.0381) with no effect on average chow meal frequency. **(M-O)** SON OT neuron silencing decreased HFHS intake at 4 h into the dark cycle (Fisher’s LSD planned comparisons 4 h: t(10.74)=2.273, p=0.0446) while having no effect on HFHS meal frequency or average HFHS meal size. (DREADDs Activation: chow n=12; sucrose n=7; HFHS n=6; within-subjects design for drug treatments; TTLC Silencing: CON n=10; SON Silenced n=7, between-subjects for group; Data are means ± SEM; *p<0.05, **p<0.01, ***p<0.001).

Chronic silencing of synaptic transmission from SON OT neurons (Figure 1C-D) significantly decreased average chow intake at 0.5 h and 1 h into the nocturnal cycle (Figure 3J), an effect driven by a decrease in 0.5 h average chow meal size (Figure 3L). There was no effect of SON OT neuron silencing on chow meal frequency (Figure 3K). When maintained on HFHS, chronic silencing of OT neurons in the SON reduced average cumulative HFHS intake at 4 h into the nocturnal cycle (Figure 3M) while having no effect on HFHS meal frequency (Figure 3N) or average HFHS meal size (Figure 3O). There were also no significant differences in SON silenced animals vs. controls for bodyweight (Supplemental Figure 1A) or change in body weight since surgery (Supplemental Figure 1B). Taken together with results from excitatory DREADDs experiments above, these findings suggest that the endogenous role of SON OT neurons involves increasing food intake by increasing meal size, with differing effects depending on macronutrient composition of the meal.

### Simultaneous activation of both PVH and SON OT neuron populations has no effect on food intake

To investigate the interconnected roles of each OT neuron population we used the same excitatory DREADDs approach to simultaneously activate both the PVH and SON populations. Co-activation of PVH and SON OT neuron populations had no effect on cumulative chow intake (Figure 4A) or average chow meal size (Figure 4C). Activation of both populations transiently increased chow meal frequency at 2 h (Figure 4B) but otherwise had no effect on consumption patterns. There was no effect of co-activation on cumulative HFHS intake, HFHS meal frequency, or average HFHS meal size (Figure 4D-F). Similarly, there was no effect of activation of both PVH and SON populations on cumulative liquid sucrose intake, liquid sucrose meal frequency, or average liquid sucrose meal size (Figure 4G-I).

**Figure 4.**
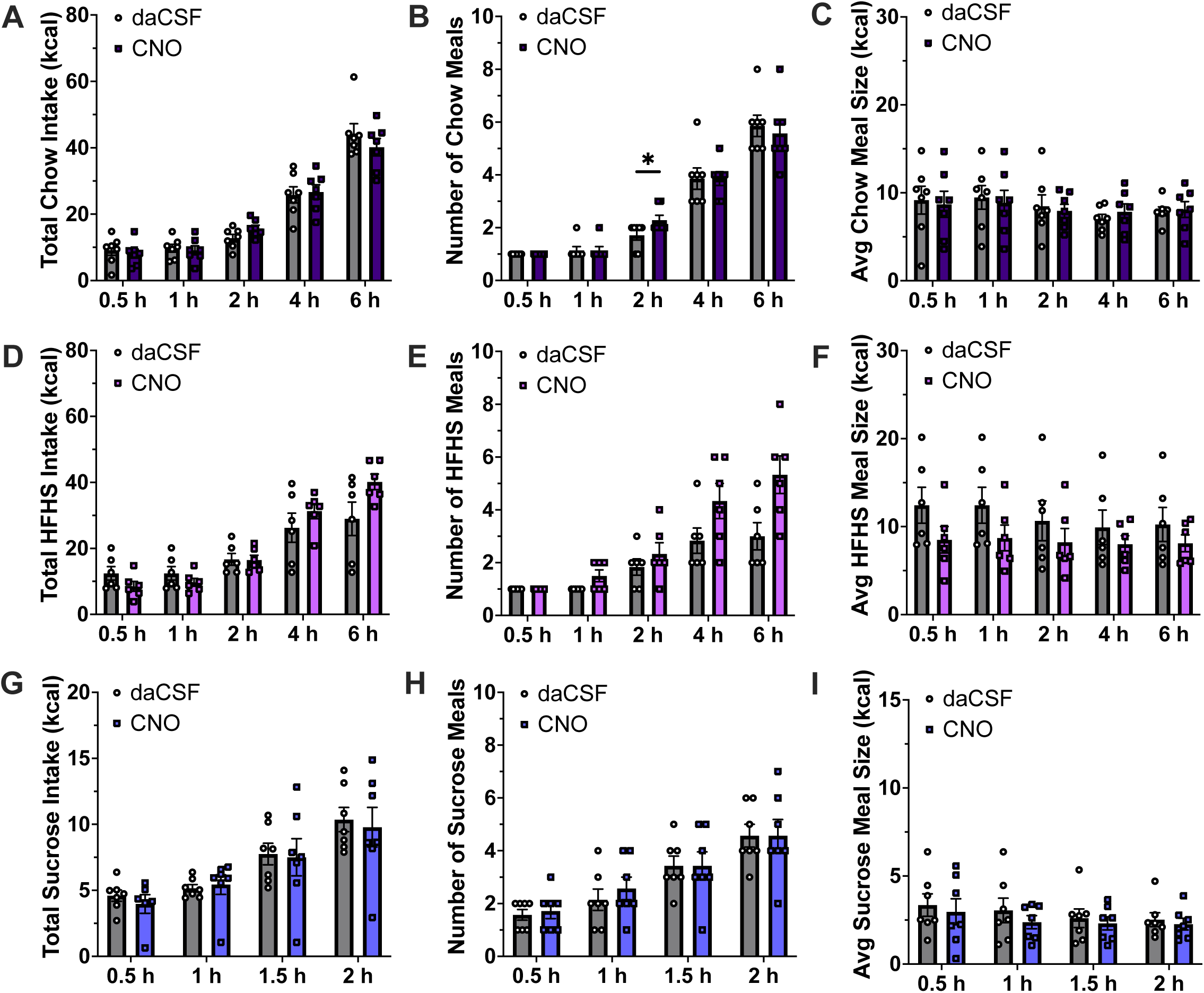
Simultaneous chemogenetic activation of both PVH and SON OT neuron populations has no effect on food intake. **(A-C)** Activation of both PVH and SON OT neurons via ICV CNO had no effect on cumulative chow intake or average chow meal size though it did transiently increase chow meal frequency at 4 h (t(6)=2.828, p=0.03). **(D-F)** Activation of PVH + SON OT neurons had no effect on cumulative HFHS intake, HFHS meal frequency or average HFHS meal size. **(G-I)** There was no effect of PVH + SON OT neuron activation on cumulative sucrose intake, liquid sucrose meal frequency or average liquid sucrose meal size. (chow n=7; sucrose n=7; HFHS n=7; all within-subjects design for drug treatments; Data are means ± SEM; *p<0.05).

As a control to eliminate the possibility of CNO administration itself playing a role in altering food intake independent of functional OTp AAV (e.g., DREADDs or TTLC), another group of animals received a nonfunctional control OTp Venus virus in both the PVH and SON regions and then had CNO infused ICV on test days. Administration of CNO in this control group had no significant effect on intake, meal frequency, or average meal size for either chow, sucrose, or HFHS conditions (Supplemental Figure 2A-I).

## DISCUSSION

Exogenous administration of OT either centrally or peripherally reduces food intake in both rodents and humans [7-9; 11; 36]. However, OT pharmacological studies fail to mimic physiological OT release patterns from OT-producing neurons in the hypothalamus. Further, the extent to which OT neuron subpopulations in the PVH and SON differentially impact feeding behavior is unknown. Here we sought to explore the individual contributions of these different OT neuron subpopulations on modulating food intake control using site-specific viral-mediated chemogenetic activation and synaptic silencing approaches [30]. Results revealed that PVH OT neuron activation significantly decreased food intake on a standard chow diet through a reduction in meal size, while having no significant effect on consumption of liquid sucrose or at high-fat sucrose-enriched palatable diet (HFHS). Consistent with these gain-of-function results, chronic silencing of synaptic transmission from PVH OT neurons resulted in an increase in HFHS intake during the early nocturnal feeding period through an increase in average meal size. Taken together, these findings support a role for PVH OT neurons in reducing food intake by enhancing the efficacy of within-meal vagus nerve-mediated satiation signals to terminate a meal. These findings agree with previous pharmacological studies supporting an anorexigenic role for OT signaling, especially in the hindbrain [7; 13]. Recent OT neuron connection mapping analyses in rodents show that PVH OT neurons have dense projections to the hindbrain, including to the dorsal vagal complex (DVC) comprised of the nucleus tractus solitarius (NTS), the area postrema (AP), and the dorsal motor nucleus of the vagus nerve (DMX) [20]. DVC neurons, particularly in the NTS, express OTRs that have been shown to decrease food intake when activated by direct OT administration [12; 13; 22]. Thus, our collective results from targeting PVH OT neurons are consistent with previous pharmacological and neuroanatomical studies that suggest that PVH OT neurons amplify satiation signals to decrease meal size via DVC projections.

Surprisingly, activation of SON OT neurons resulted in a significant increase in chow intake via elevated meal size. SON OT neuron activation also increased consumption of liquid sucrose, although these results were mediated by an increase in meal frequency. In support of these findings, SON OT neuron silencing significantly reduced chow intake and meal size during the early nocturnal feeding period, indicating that SON OT neurons may play an endogenous role in promoting food intake by enhancing meal size during periods of active eating. While these finding differs from most previous studies that solely identify an anorexigenic role for OTR signaling [7; 18], recent results from our lab reveal that OT administration into the dorsal subregion of the hippocampus, a region critical for both food intake control and social memory, can increase food intake under conditions where food is consumed with a familiar conspecific [37]. Further, our study identified an endogenous role for hippocampus OTR signaling in promoting prosocial eating, as targeted hippocampal OTR viral-mediated knockdown eliminated the social facilitation of eating effect [37]. Future studies are needed to determine whether SON vs. PVH OT neurons differentially engage the hippocampus, and whether these subpopulations differ with regards to mediating interactions between food intake and social factors. Given that very few or no OT-immunoreactive axons are not present in the dorsal CA1 hippocampus [20; 30; 38], it is likely that OT neurons engage hippocampal OTRs through volume transmission – that is, humoral-like “bulk flow” transmission of peptides through release into cerebral spinal fluid and/or the brain interstitial space [33].

The bidirectional differences in the effects of these two hypothalamic OT neuron subpopulations on caloric consumption might be explained by differences in their morphology, anatomical inputs and outputs, mechanisms of OT transmission, and/or OTR properties of downstream targets. For example, OT neurons within the PVH and SON are predominantly categorized into two main types, magnocellular and parvocellular (preautonomic) neurons, though recent studies argue that there may be four distinct types based on genetic profiles [21]. Magnocellular OT neurons from both hypothalamic nuclei innervate various forebrain areas and release OT into the blood via the posterior pituitary. They contain many large dense core vesicles (LDCV) and have been shown to release OT somato-dendritically as well [39]. While these magnocellular OT neurons are present in both subpopulations, in the PVH but not the SON there are also a small number (at least in rats) of parvocellular OT neurons that do not possess connections to the neurohypophysis, but instead connect directly to brainstem, midbrain, and forebrain targets [40]. While parvocellular OT neurons in rats are thought to make up only a small percentage of all OT neurons, their projection targets are numerous and they have been shown to act as regulators of magnocellular OT neuron activity as well [41–43]. Consistent with parvocellular OT neurons being present only in the PVH, OT PVH and SON neurons differ with regards to their projection profile [20]. One characteristic feature of the PVH is that it forms reciprocal connections sending outputs to most of the regions it receives input from, which could allow for extrahypothalamic OTR-expressing neurons to tune OT neuron activity depending on behavior context [44]. An exception to this is the hindbrain which sends relatively little input to the PVH, yet receives dense output into the DVC from PVH OT neurons [20], which as mentioned above may drive PVH OT neurons’ anorexigenic effects. Unlike PVH OT neurons, however, SON OT neurons appear not to send any outputs to the hindbrain. Thus, while the PVH and SON receive inputs from mostly similar regions, SON OT neurons have relatively fewer outputs that represent a subset of those targeted by the PVH [20]. For this reason, the SON OT neurons have been thought to predominantly function in peripheral neuroendocrine signaling. While more research is required to confirm, it may be that PVH parvocellular OT neurons reduce meal size via brainstem projections, whereas SON magnocellular OT neurons have the opposite effect, potentially mediated by peripheral OT release.

Both PVH and SON project into the lateral hypothalamic area (LHA) [20]. Contained within the LHA are neurons that produce melanin-concentrating hormone (MCH), a potent orexigenic neuropeptide [45; 46]. Interactions between OT and MCH systems have been established, with OT neurons expressing MCH receptors, as well as 60% of MCH neurons expressing OTR [47]. Studies in slice preparations have shown that OT, and selective OTR agonists result in the excitation of MCH neurons, to a greater degree than other GABAergic neurons within the LHA [48]. Meanwhile MCH injected into the PVH has been shown to increase feeding [49]. However, the extent to which OT and MCH interact to influence feeding behavior has not been directly investigated. One possibility is that these two OT neuron subpopulations target different cells within the LHA, with SON but not PVH OT neurons engaging MCH neurons to promote food intake.

In addition to differences in OT cell types and projection targets for PVH vs. SON OT neurons, the bidirectional effects on food intake could be mediated by differences in downstream OTR dynamics. While there is only one OTR, it is a G-protein coupled receptor that shows heterogeneity in that it can be associated with excitatory as well as inhibitory G-proteins depending on the cell in which it is expressed [4]. Moreover, OTRs are also expressed on a wide range of cell types including GABAergic interneurons [50], glutamatergic pyramidal cells [51], and other neuroendocrine cells [52]. This heterogeneity in OTR signaling mechanism and cell-type expression could allow for OT to mediate a wide range of effects, including in a bidirectional manner depending upon the behavioral and physiological context.

One limitation of this study is the exclusion of females. In our lab we have recently identified varying effects of OT administration on food intake depending on estrus cycle stage [26]. Additionally, there are known sex differences in feeding behavior, OT production, and central OTR expression, with estrus cycle stage altering OTR expression levels in cycling females [53–56]. For these reasons females were excluded for initial studies, though this represents an important area of follow-up investigation to confirm whether or not the differential roles of PVH and SON OT neurons in regulating feeding behavior are sex-specific. Another limitation of the current study involves the technical difficulty to separately target and distinguish roles of parvocellular vs. magnocellular OT neurons within the PVH subpopulation. Use of the OT promoter targets both cell types without the ability to select between them. The ability to differentially target OT neurons based on cell type would be informative with regards to understanding the complex dynamics of hypothalamic OT neuron signaling between subpopulations. Finally, present findings should be considered with regards to the involvement of the arginine vasopressin (AVP) system, as this neuropeptide is closely related to OT (evolutionarily diverged from a combined gene) and there has been shown to be substantial cross talk between OT and AVP systems/receptors [57; 58]. While OT binds to OTR with a higher affinity than to AVP receptors, the chemogenetic activation to potentially supraphysiological levels could result in off-target effects in the AVP system [59]. The use of a selective OTR antagonist in future studies combined with the OTp chemogenetic approach could help to ensure these effects are indeed mediated by the OTR. In conclusion, our results demonstrate that separate hypothalamic OT neuron populations in the PVH and SON exert opposing influences on food intake control, serving to decrease and increase meal size, respectively. These effects are likely mediated by differences in projection targets of these subpopulations, as well as by different release dynamics mediated by different cell types (e.g., parvocellular OT neurons are exclusively in PVH). PVH OT neurons likely engage classic anorexigenic pathways in the hindbrain while SON OT neurons may have connections with orexigenic systems in the LHA MCH system, and potentially may promote food intake via unknown orexigenic peripheral signaling pathways. Overall, the current results should be taken into consideration with regards to OT-based targets for clinical obesity treatment, as OT’s endogenous role in regulating feeding behavior may be more complex than previously considered to be.

## Acknowledgements

The authors would like to thank the Kanoski Lab undergraduate research assistants for their support with the behavioral experiments as well as all funding sources.

## Funding

This work was supported by the following grants:

National Institute of Diabetes and Digestive and Kidney Diseases: DK118402, and DK104897 (to S.E.K.), F31DK137484 (to J.J.R.)

Postdoctoral Ruth L. Kirschstein National Research Service Award from the National Institute on Aging F32AG077932 (to A.M.R.H.)

Quebec Research Funds postdoctoral fellowship 315201 (LDS)

Alzheimer’s Association Research Fellowship to Promote Diversity AARFD-22-972811 (LDS)

The Synergy European Research Council (ERC) grant 101071777, SFB Consortium 1158-3, and German-Israeli Project cooperation (DIP) GR3619-1 to V.G.

---

**Supplemental Figure 1.**
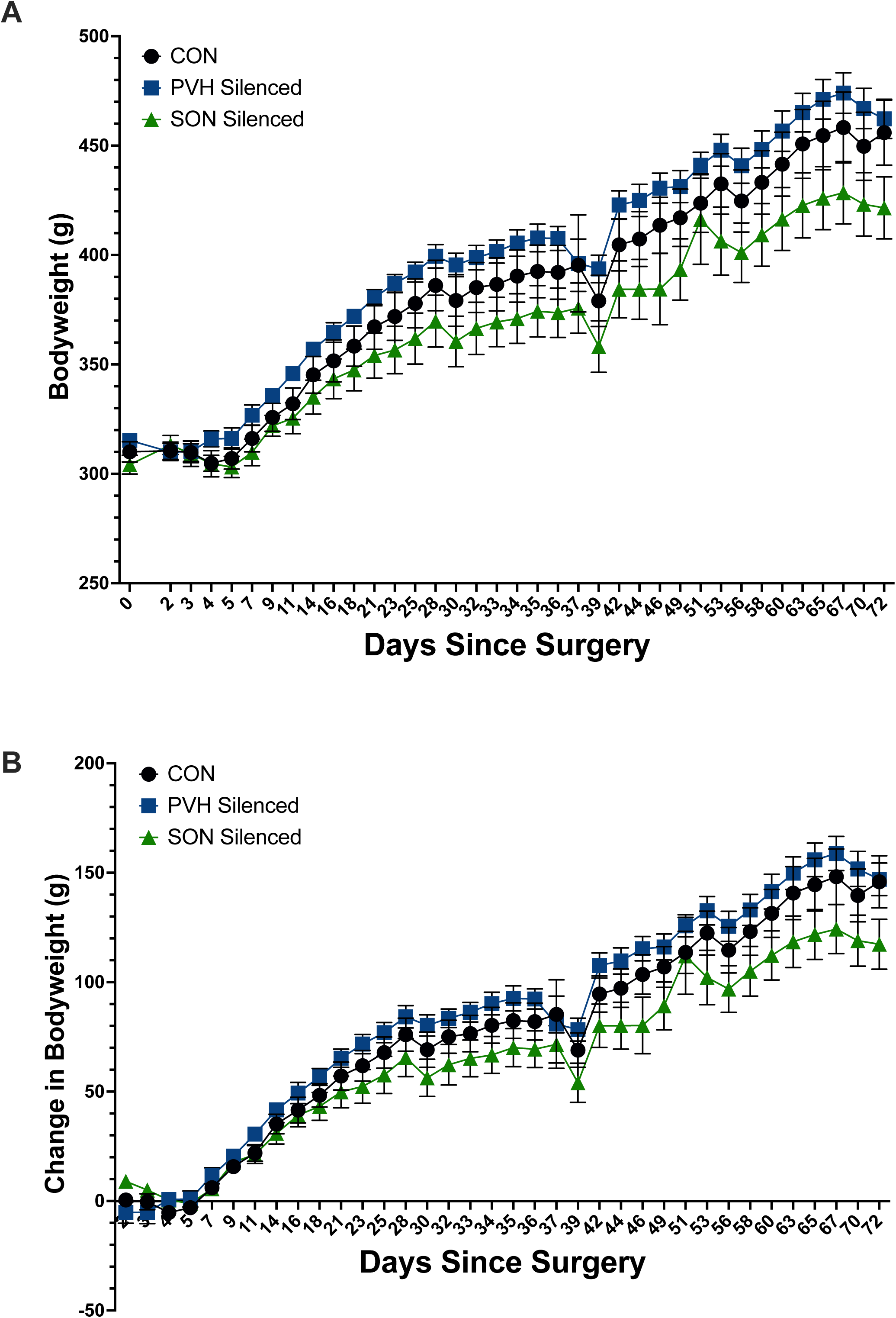
Changes in bodyweight following the expression of OTp TTLC to chronically silence synaptic transmission from OT neurons. **(A)** There were no differences in average bodyweights between groups following the expression of OTp in either the PVH or SON. **(B)** No differences were seen in the average change in bodyweight following chronic synaptic silencing of OT neurons in either the PVH or SON compared to controls.

**Supplemental Figure 2.**
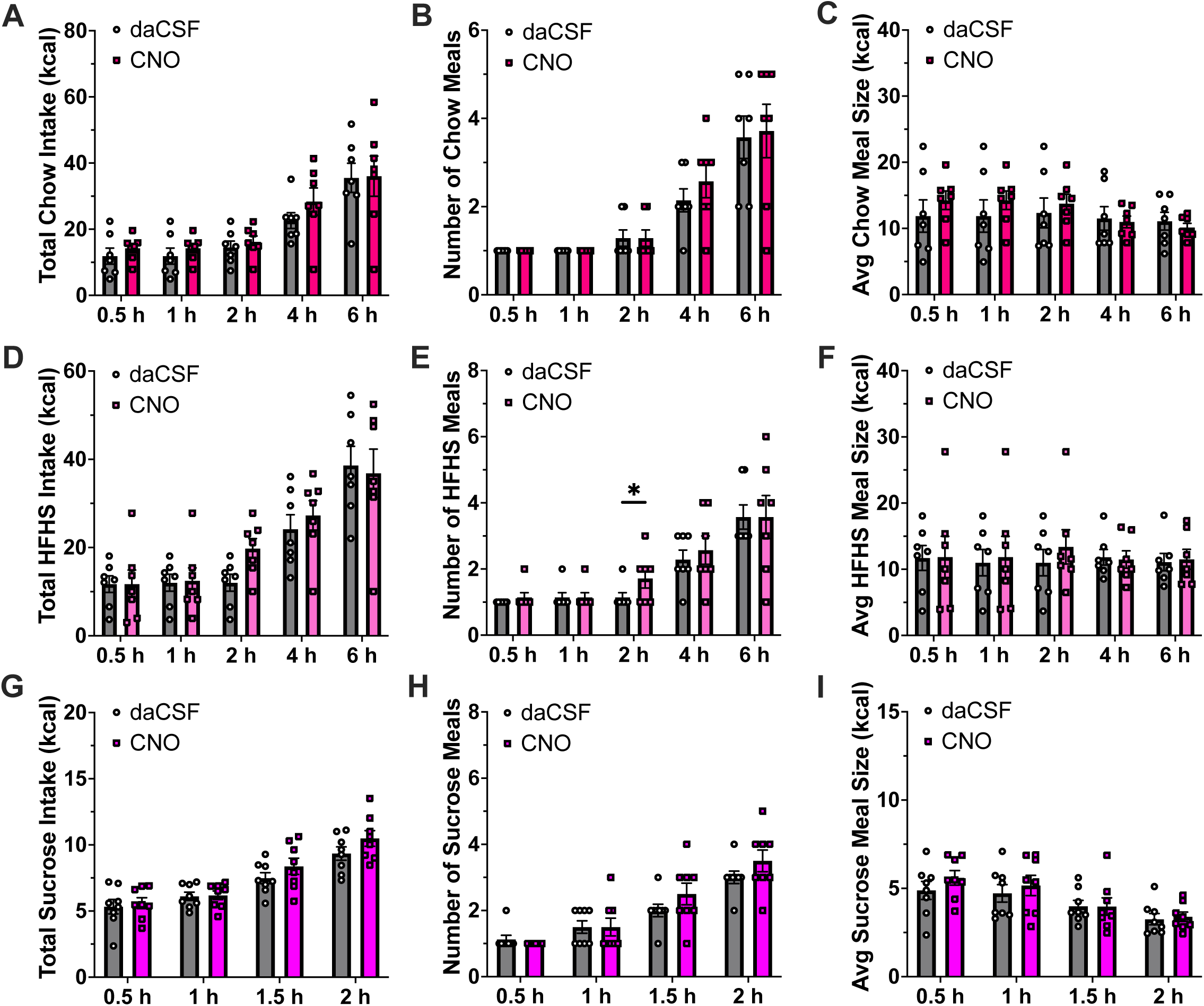
Lateral ventricle infusion of CNO has no effects on food intake. **(A-C)** ICV administration of CNO in animals with control virus (anterograde tracer, not DREADDs) had no effect on cumulative chow intake, chow meal frequency or average chow meal size. **(D-F)** ICV administration of CNO had no effect on cumulative HFHS intake or average HFHS meal size, though there was a transient increase in HFHS meal frequency at 2 hr (t(6)=2.828, p=0.03). **(G-I)** CNO administration in control animals had no effect on cumulative sucrose intake, liquid sucrose meal frequency or average liquid sucrose meal size. (CNO Control: chow=7, sucrose=8, HFHS=7; all within-subjects design for drug treatments; Data are means ± SEM; *p<0.05).

